# Rapid Detection of Carbapenemase-Producing Organisms Using a Luminescent Biosensor

**DOI:** 10.1101/2025.04.01.646590

**Authors:** Mitchell A. Jeffs, Shreyas Bhat, Josephine L. Liu, Rachel A. V. Gray, Xena X. Li, Prameet M. Sheth, Christopher T. Lohans

## Abstract

Carbapenemase-producing organisms (CPOs) pose an urgent global health threat due to their ability to inactivate carbapenems, a group of last resort antibiotics. Infections caused by these pathogens are associated with poor patient outcomes, high mortality rates, and added burden to infection prevention and control programs, making early detection vital to ensure optimal antimicrobial therapy and appropriate implementation of infection control practices. In this study, we report the application of a luminescent whole-cell biosensor for the rapid detection of CPOs. This biosensor provides positive test results within 2.5 hours, inclusive of setup time, and has been validated with a panel of laboratory and clinical isolates producing a diverse range of carbapenemases (KPC, NDM, IMP, VIM and OXA-48-like). The assay successfully identified all CPO isolates tested with a sensitivity of 100%, including strains producing weak OXA-48-like carbapenemases which are sometimes missed by currently used detection methods. The assay also demonstrated a specificity of 100 %, with all non-CPO clinical isolates testing negative under the optimized assay conditions. Due to the rapid time-to-positivity, minimal setup requirements, and high sensitivity, this test could serve as an attractive alternative to CPO detection methods currently employed by clinical microbiology labs, and could also facilitate CPO screening in other settings (*e.g.,* environmental, agricultural).

## INTRODUCTION

β-lactams are currently the most commonly prescribed class of antibiotics worldwide, and include the penicillins, cephalosporins, carbapenems and monobactams.^1^ Several drugs belonging to this class (*e.g.,* carbapenems such as meropenem, ertapenem and imipenem) are considered last-resort antibiotics for targeting multi-drug resistant pathogens. The carbapenems are used for a wide array of clinical indications including urinary tract infections, pneumonia, intra-abdominal infections and meningitis, among others.^2^

The global emergence and increasing prevalence of carbapenemase-producing organisms (CPOs) poses a massive global health concern, leading to their classification as a critical group of pathogens by the WHO.^3^ Carbapenemases confer bacteria with resistance against carbapenems (as well as other types of β-lactams) by hydrolyzing the β-lactam ring in the core structure of these antibiotics. Carbapenemases are a subset of the β-lactamases, a diverse group of hydrolytic enzymes which also include the penicillinases and extended-spectrum β-lactamases (ESBLs). These enzymes can be further subdivided into several classes based on the Ambler classification system: class A (*e.g.,* KPC) and class D (*e.g.,* OXA) enzymes rely on an active site serine residue to catalyze carbapenem hydrolysis, while class B (*e.g.,* NDM, IMP, VIM) carbapenemases utilize zinc ions to catalyze hydrolysis. These enzymes are produced by a variety of bacterial pathogens, including *Escherichia coli*, *Pseudomonas aeruginosa*, *Acinetobacter baumannii*, *Klebsiella pneumoniae* and *Enterobacter cloacae*; infections caused by these CPOs are associated with high mortality rates.^4–6^ Thus, rapid detection of CPOs in clinical settings is vital to ensure administration of appropriate antimicrobial therapy, as well as the prevention and containment of outbreaks caused by these pathogens.

Several different assays are currently employed for the detection of CPOs. Traditional phenotypic tests endorsed by the Clinical and Laboratory Standards Institute (CLSI) include growth-based assays such as the modified carbapenem inactivation method (mCIM). Although the mCIM is widely employed in hospital settings due to its low cost per test and ease of use, this assay suffers from long turn-around times, typically requiring 18 – 24 h (not including the time needed for the initial CPO isolation and culturing) before results are obtained.^7,8^ Other, more rapid assays have also been developed which test the phenotypic properties of a potential CPO. Currently the sole rapid carbapenemase activity assay endorsed by the CLSI is the colorimetric CARBA NP test, which has a turnaround time of approximately 2 hours.^9,10^ Despite high sensitivity for the detection of KPC-, NDM-, IMP- and VIM-type carbapenemases, some labs have observed that this assay suffers from lower sensitivity for CPOs that produce OXA-type class D carbapenemases.^11–15^ Other rapid tests employ fluorogenic or chromogenic β-lactams,^16–18^ but their use may be limited due to costs associated with the synthesis or purchase of these non-clinical substrates. Lateral flow assays such as the NG-Test Carba-5 have also been developed, which provide extremely rapid results (15 - 30 min) and demonstrate a high degree of sensitivity and specificity for the “big five” carbapenemases (OXA-48, KPC, IMP, VIM, NDM).^19^ However, this assay cannot detect carbapenemases which do not belong to one of these subclasses, and the cost per test is high. In recent years, the use of genotypic tests has expanded due to their high sensitivity and quick turn-around times. Several polymerase chain reaction (PCR) panels have been applied to the detection of carbapenemase genes in potential CPOs.^20–23^ Despite these advances, genotypic assays are less feasible in lower resource settings due to high costs associated with equipment and reagents, the need for highly trained staff,^24^ and potential limitations associated with the detection of uncommon and novel carbapenemase variants.

In this study, we report the development of a luminescence assay for the rapid detection of CPOs. This test employs a biosensor developed by our lab to study β-lactamase inhibition,^25^ with major modifications to optimize the assay for carbapenemase detection. We prepared a plasmid in which transcription of a luminescent reporter is under the control of the carbapenem- inducible *ampC* promoter. This plasmid is maintained in a laboratory strain of *E. coli* which serves as a whole-cell biosensor. Exposure of biosensor cells to a carbapenem promotes the formation of peptidoglycan fragments, ultimately resulting in the production of a luminescent signal (Figure 1A). However, upon addition of a CPO to the assay media, the carbapenem is degraded, reducing the luminescent signal. The workflow of this assay begins with the addition of colonies of the test strain to a lysis solution, which is then incubated with imipenem, then transferred to a suspension of biosensor cells (Figure 1B). Luminescence values are recorded after a 90 min incubation period.

**Figure 1.**
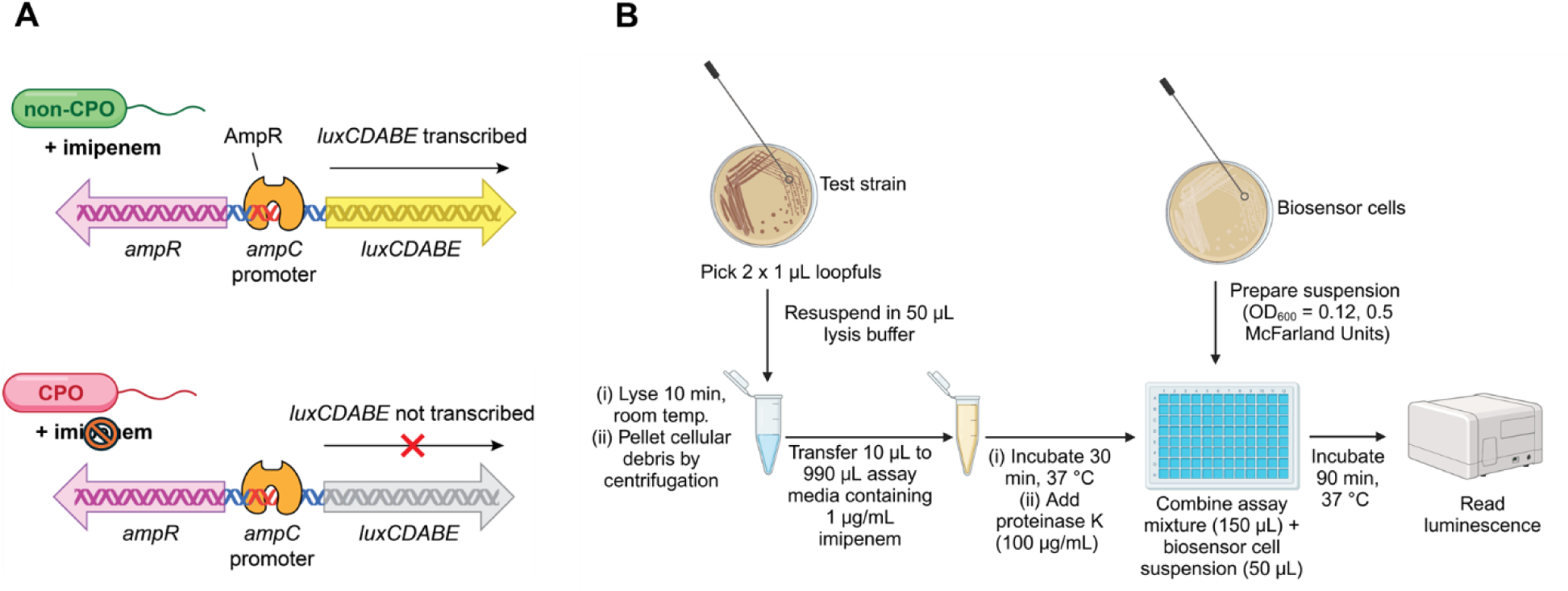
(A) Rationale for the developed biosensor. In samples which do not contain a CPO, the carbapenem imipenem is not hydrolyzed, resulting in strong luminescence production by the biosensor. In contrast, imipenem is degraded in samples which contain a CPO, resulting in low luminescence production by the biosensor. **(B)** Graphical overview of the assay workflow.

β-lactamase inhibitors (*e.g.,* avibactam, vaborbactam) are often administered alongside β- lactam antibiotics, protecting them from β-lactamase-catalyzed degradation. However, carbapenemase variants which are less susceptible to inhibition have begun to emerge in recent years.^26–28^ Current rapid testing methods do not typically incorporate clinically used inhibitors into their workflow; instead, labs tend to rely on growth-based susceptibility testing methods to evaluate inhibitor efficacy.^29,30^ These tests suffer from similar limitations as the mCIM, requiring overnight culturing (18 - 24 h) before results can be interpreted. As resistance to these β-lactamase inhibitors increases in prevalence, there is a need for rapid CPO detection methods that can also inform on inhibitor susceptibility. We show that carbapenemase inhibitors such as avibactam can be easily incorporated into the biosensor assay, allowing their efficacy against the carbapenemase(s) produced by a CPO to be evaluated.

Our CPO detection assay was validated with laboratory *E. coli* strains producing common carbapenemases (Table S1) (*i.e.,* KPC-2, NDM-1, IMP-1, and OXA-48), then tested against a panel of clinical CPO and non-CPO isolates (Tables S2 and S3). This panel consisted of a diverse range of bacterial species and carbapenemase types (where applicable), representative of what clinical microbiology labs commonly encounter. The assay provides positive results within 2.5 hours, inclusive of setup, and displays excellent sensitivity (100%) and specificity (100 %). Thus, this biosensor assay could serve as a valuable tool in clinical microbiology labs for the rapid detection of CPOs, ensuring that infection prevention and control (IPAC) measures (*e.g.,* contact precautions, patient isolation, increased room sanitation)^31^ can be implemented or de-escalated more quickly. In addition to clinical applications, this assay could be used as a primary screening tool for CPO detection in environmental (*e.g.,* wastewater) and agricultural samples. Surveillance of CPOs in these settings is key, as agricultural and environmental sources of CPOs may act as reservoirs for colonization/infection and can promote the spread of carbapenemase-encoding genes through horizontal gene transfer.^32–34^

## RESULTS AND DISCUSSION

The assay described in this study employs a carbapenem-inducible whole-cell biosensor which, following exposure to carbapenem antibiotics, produces increased levels of 1,6- anhydromuropeptides leading to luminescence production.^25^ These anhydromuropeptides are produced as a result of β-lactam-induced damage to peptidoglycan in the bacterial cell wall following inhibition of penicillin-binding protein 4 (PBP4).^35,36^ However, if the carbapenem in the assay media is first degraded by a carbapenemase, the luminescent signal of the biosensor cells is significantly reduced.^25^ Thus, we proposed that our biosensor platform could be adapted and applied to the rapid detection of CPOs (Figure 1A, B).

Preliminary testing of the biosensor was conducted with a library of *E. coli* BW25113 strains producing some of the most prevalent plasmid-encoded carbapenemases (*i.e.,* KPC-2, NDM-1, IMP-1, and OXA-48) (Figure 2A, Table S1). In these preliminary tests, colonies of CPO lab strains grown on CAMHB agar were first suspended in assay media. Incubation of these cell suspensions with the carbapenem imipenem (1 µg/mL) and *E. coli* BW25113 pAMPLUX biosensor cells for 90 minutes yielded high detection scores (> 8) for all carbapenemases tested (Figure 2A). These detection scores were determined by dividing the luminescent readings for a carbapenemase-negative sample with the luminescent readings for the test samples. Notably, even the strain producing the carbapenemase OXA-48 tested strongly positive, with detection scores comparable to the other carbapenemases tested. OXA-48 enzymes are notoriously difficult to detect due to their relatively weak hydrolytic activity against carbapenems, when compared to other types of carbapenemases.^37,38^ The difficulties associated with detecting OXA-48-like enzymes is concerning, as these enzymes have greatly increased in prevalence worldwide.^39^ We also tested a series of carbapenemase-producing *E. coli* lab strains which lack OmpF, a key porin used by β-lactam antibiotics to enter the cell.^40,41^ All Δ*ompF* strains producing carbapenemases still tested strongly positive (Figure S1).

**Figure 2:**
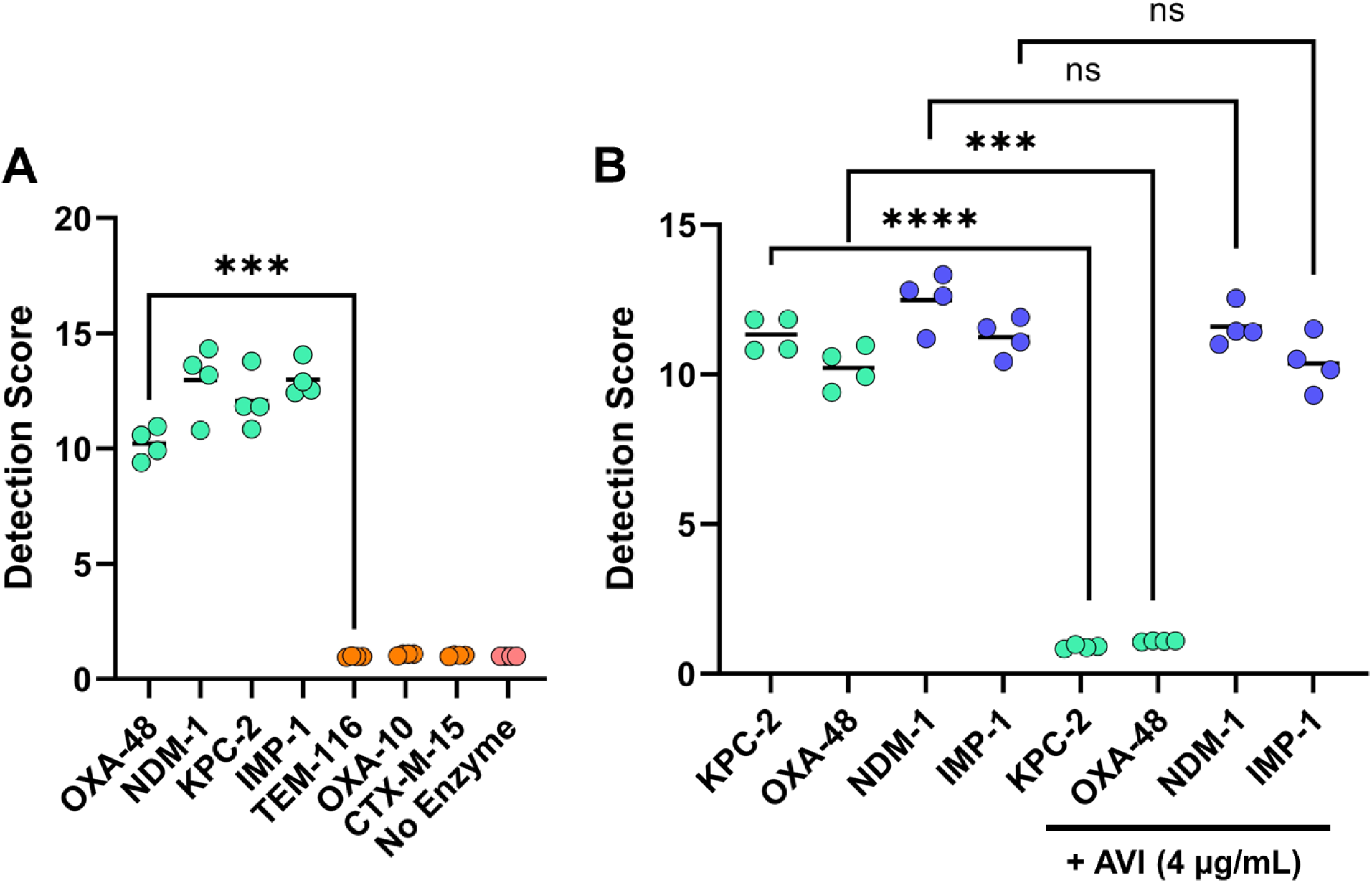
Validation of the biosensor-based CPO detection assay with carbapenemase- producing *E. coli* lab strains. **(A)** Detection scores for *E. coli* producing carbapenemases OXA- 48, NDM-1, IMP-1, and KPC-2 (light blue circles), penicillinases TEM-116 and OXA-10 (orange circles), the ESBL CTX-M-15 (orange circles) or no β-lactamase (pink circles). **(B)** Comparison of detection scores for *E. coli* strains producing either serine β-lactamases (light blue circles) or metallo-β-lactamases (dark blue circles) in tests using imipenem alone versus tests combining imipenem with 4 µg/mL avibactam (AVI). A reduction in detection score in the presence of avibactam indicates that carbapenemase activity has been inhibited. Detection scores were determined by dividing the luminescent readings for a carbapenemase-negative sample with the luminescent readings for the test samples. All strains were tested in quadruplicate. ns = nonsignificant; ***: p < 0.001; ****: p < 0.0001.

*E. coli* BW25113 strains expressing plasmid-encoded penicillinases and ESBLs (TEM- 116, CTX-M-15, OXA-10) were used to assess the specificity of the assay. These β-lactamases cannot degrade imipenem,^42^ and therefore should not cause a reduction in luminescence in the assay. As expected, the TEM-116, CTX-M-15 and OXA-10-producing strains all yielded low detection scores comparable to that obtained for a β-lactamase-negative control strain (Figure 2A).

We next tested whether the biosensor assay could be employed to evaluate the efficacy of β-lactamase inhibitors against the carbapenemase produced by a CPO. To the assay, we added a second treatment containing a fixed concentration of the β-lactamase inhibitor avibactam (4 µg/mL) in combination with imipenem and compared the resulting detection scores to those for samples treated with imipenem alone (Figure 2B). KPC-2- and OXA-48-producing *E. coli* BW25113 strains demonstrated notable reductions in detection scores following the combined imipenem and avibactam treatment compared to imipenem alone. In contrast, NDM-1- and IMP- 1-producing *E. coli* strains did not exhibit a significant reduction in score, indicating that avibactam did not prevent the hydrolysis of imipenem by these CPOs. These results are consistent with the inhibitory mechanism of avibactam, which targets many serine β-lactamases like KPC-2 and OXA-48, but not metallo-β-lactamases like NDM-1 and IMP-1.^43^

With the promising results obtained in tests employing laboratory CPO strains, we proceeded to evaluate the efficacy of the assay when applied to the detection of clinical CPO isolates. However, during preliminary testing, the detection scores obtained for several clinical strains were markedly lower than those for the model laboratory CPO strains (data not shown). This likely reflects reduced outer membrane permeability or lower levels of carbapenemase production compared to the lab strains.

To address the potential impact of cell wall permeability on detection, an additional lysis step was introduced to the assay workflow. Prior to testing clinical isolates, the concentration of the BugBuster lysis solution was first optimized such that the test strain was successfully lysed while also ensuring minimal downstream impact on the biosensor cells. The lysis of *E. coli* BW25113 cells was observed to be most effective at BugBuster concentrations greater than 25 % v/v (Figure S2). However, 50 % and 100 % v/v BugBuster solutions had a significant impact on luminescence production by the biosensor cells, while the 25 % solution did not (Figure S3). Thus, a solution containing 30% v/v BugBuster was used to ensure adequate lysis while also minimizing the impact on biosensor cells.

To overcome the possibility of low carbapenemase expression or weak carbapenemase activity, we introduced a 30 min pre-incubation step in which test lysates were incubated with imipenem prior to the addition of biosensor cells. Several clinical isolates that produce OXA-48- like enzymes, which exhibit relatively weak carbapenemase activity, were used to evaluate the impact of the pre-incubation on detection. The results show that pre-incubating cell lysates with imipenem prior to the addition of biosensor cells resulted in markedly increased detection scores for isolates that tested only weakly positive without pre-incubation (Figure 3).

**Figure 3:**
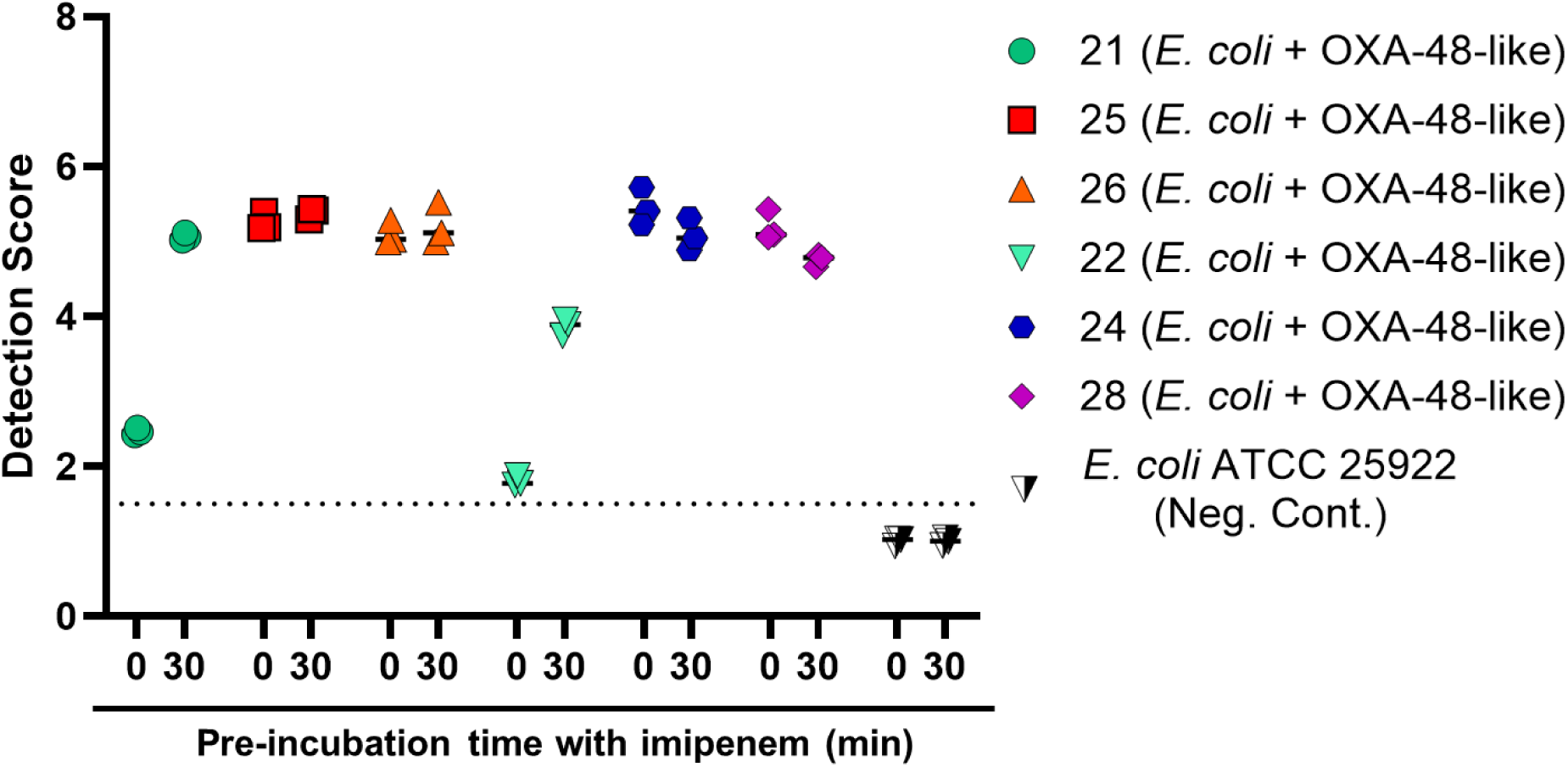
Impact of pre-incubating clinical OXA-48-like-producing *E. coli* isolates with imipenem. The detection scores for selected OXA-48-like-producing *E. coli* lysates (clinical isolates 21, 22, 24-26, 28; Table S2) incubated with 1 µg/mL imipenem for 30 min prior to the addition to biosensor cells, compared to samples for which imipenem and biosensor cells were added at the same time. The 30 min pre-incubation time improved the detection of isolates which exhibited low detection scores with no pre-incubation. The horizontal dotted line indicates the cutoff detection score to define a positive vs. negative result (samples which yielded detection scores greater than 1.5 were defined as positive). All isolates were tested in triplicate.

With this optimization completed, we proceeded to test the sensitivity of the biosensor assay against a panel of 50 clinical CPO isolates (Tables 1, S2). The overall sensitivity of the assay was 100%, with all isolates producing NDM (n = 12/12), VIM (n = 14/14), KPC (n = 15/15), KPC + NDM (n = 1/1) and OXA-48-like (n = 8/8) enzymes testing positive. Samples with a detection score greater than 1.5 were defined as positive; this value was chosen as it lies well above the average error on luminescence readings (approx. 5-10 %), thus ensuring that CPO-positive and negative samples can be distinguished. We determined the imipenem MICs for all isolates (Table S2), and compared these MICs to the biosensor detection scores (Figure 4). The detection scores remained consistent across all isolates with MICs ≥ 4 µg/mL, with an average detection score of 5.9. While a reduction in the average detection score to 4.3 was observed for isolates with low imipenem MICs (≤ 2 µg/mL), these strains could still be easily distinguished from non-CPO strains.

**Table 1:** Summary of imipenem MICs and biosensor detection results for the clinical CPO isolates.

| Carbapenemase Ambler Class (n) | Carbapenemase Subtype (n) | Imipenem MIC ($\mu\text{g/mL}$) | | | | | Biosensor Result* |
| --- | --- | --- | --- | --- | --- | --- | --- |
| | | $\geq 32$ | 16 | 8 | 4 | $\leq 2$ | Positive |
| Class A (15) | KPC (15) | 1 | 0 | 6 | 7 | 1 | 15/15 |
| Class B (26) | VIM (14) | 3 | 5 | 3 | 3 | 0 | 14/14 |
|  | NDM (12) | 4 | 3 | 4 | 1 | 0 | 12/12 |
| Class A + B (1) | KPC + NDM (1) | 0 | 1 | 0 | 0 | 0 | 1/1 |
| Class D (8) | OXA-48-like (8) | 2 | 3 | 0 | 0 | 3 | 8/8 |
| Total (50) |  |  |  |  |  |  | 50/50 |

\* Samples which yielded detection scores greater than 1.5 were defined as positive

**Figure 4:**
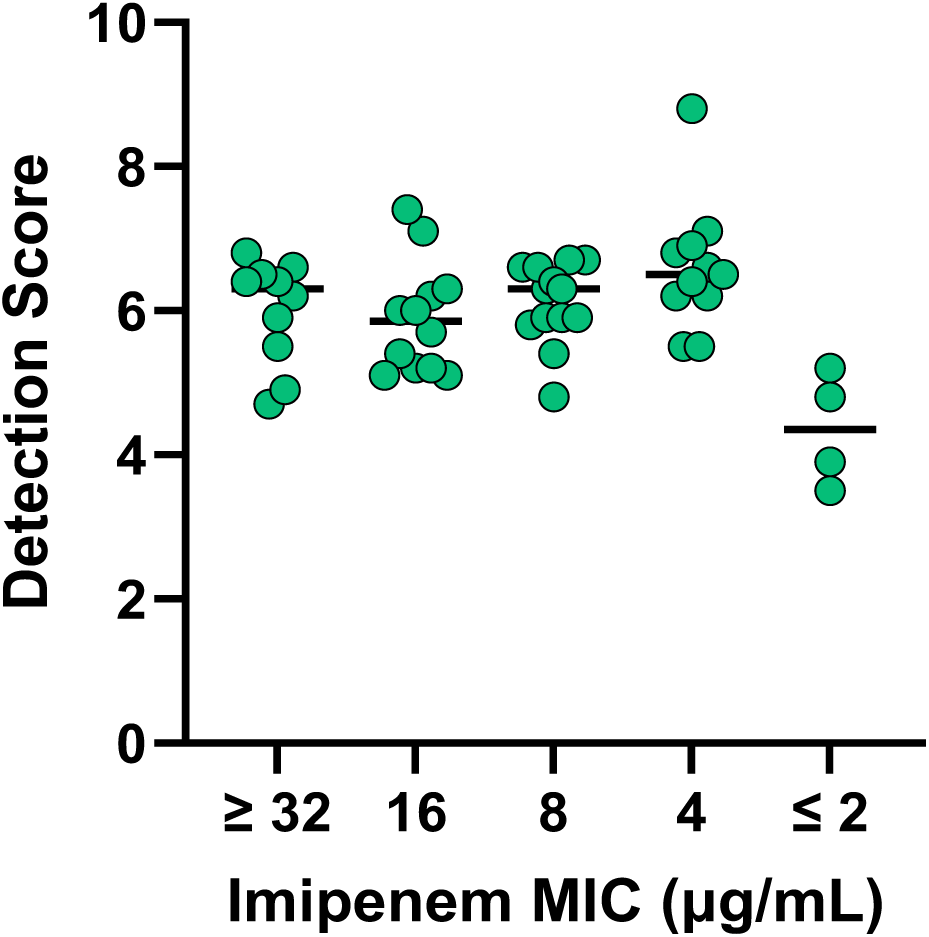
D**i**stribution **of detection scores for CPO isolates, grouped according to their imipenem MICs.** Detection scores remain consistent across the imipenem MIC brackets ≥ 4 µg/mL, with average detection scores > 5.9. For isolates with low imipenem MICs (≤ 2 µg/mL), the average detection score is reduced to 4.3.

The widely used CARBA-NP test can be used to detect CPOs in approx. 2 hours, and exhibits a high degree of sensitivity for the detection of bacteria that produce KPC, NDM, IMP and VIM-type carbapenemases.^9,11^ However, some groups have reported that the CARBA-NP test exhibits lower sensitivity for isolates producing carbapenemases with weaker hydrolytic activities. A study by Tijet *et al.* reported that 5/5 GES-5-producing CPOs and 31/39 OXA-48-like-producing CPOs falsely tested negative,^11^ and Istar *et al*. observed that the CARBA-NP test exhibited a low degree of sensitivity among OXA-48-producing-Enterobacterales.^12^ With these limitations in mind, we compared the efficacy of the CARBA-NP test to our whole-cell biosensor assay for the detection of OXA-48-like-producing isolates from our CPO library. Overall, the CARBA-NP yielded positive results for 6/8 OXA-48-like-producing *E. coli* isolates, with two isolates yielding a false-negative result (Figure S4). In contrast, our biosensor assay successfully detected 8/8 OXA- 48-like-producers. The average detection score for these isolates was 4.8, which well exceeds the positivity threshold of 1.5. Two strains that exhibited imipenem MICs of 2 µg/mL, which only slightly exceeds the CLSI susceptibility breakpoint of 1 µg/mL, still tested strongly positive with detection scores of 3.5 and 3.9. It should be noted that only a small library of OXA-48-like-producing isolates was tested in this study, and the ability of the biosensor assay to detect CPOs producing OXA-48 and other weak carbapenemases should be further validated with a larger library of clinical isolates.

To verify the specificity of the biosensor assay, we tested a library of ESBL-producing Enterobacterales isolates which did not flag positive for carbapenemase production. In the first round of testing, while 26/29 isolates tested negative, three isolates unexpectedly yielded high detection scores (Table S3). Note that one isolate (N3) did not yield sufficient growth on CAMHB- ampicillin plates for testing and was excluded from the study. We proceeded to perform follow-up testing on the three ESBL-producing *E. coli* isolates (N4, N23, N24) which gave positive results. UV-Vis spectrophotometric β-lactamase assays indicated that N4, N23 and N24 did not hydrolyze imipenem, consistent with the expected absence of carbapenemase production by these isolates (Figure S5). Interestingly, the luminescence of the biosensor cells decreased following treatment with lysed N4, N23 and N24 cells in the absence of imipenem (Figure S6), suggesting that these strains may be generating a product that directly targets the biosensor cells.

Like other Enterobacterales, many *E. coli* strains produce bacteriocins, antibacterial proteins which can kill other *E. coli* strains.^44–46^ As such, the reduction of biosensor luminescence following exposure to N4, N23 and N24 cell lysates could indicate the presence of bacteriocins. With this in mind, we incorporated the broad-spectrum protease proteinase K (PK) into the assay workflow to test whether it could overcome the false positives associated with these isolates. Addition of a PK treatment after the imipenem pre-incubation step successfully reduced the detection scores for N4, N23 and N24 to levels consistent with other CPO-negative isolates (Tables 2, S3). These results suggest that the production of bacteriocins or other antimicrobial proteins by *E. coli* isolates N4, N23 and N24 was indeed responsible for the high detection scores, and that the addition of PK to the workflow can overcome this hurdle. All 29 CPO-negative strains were re- tested with the PK incubation step, resulting in no false positives and a specificity of 100 % (Tables 2, S3). We also verified that the addition of PK did not impact detection of CPOs, with comparable detection scores being obtained for representative NDM-, KPC- and OXA-48-like-producing CPOs (Table S4). Notably, the addition of PK reduced a markedly high detection score for a VIM-producing *E. cloacae* isolate (# 30) to a score on-par with other CPOs, which suggests that this isolate may also have been producing an antimicrobial protein. Although the addition of PK will address the production of most antimicrobial proteins or peptides, it may not account for antibacterial agents that are not susceptible to degradation by PK.

**Table 2:** Summary of biosensor performance for non-CPO, ESBL-producing clinical isolates with and without proteinase K (PK; 100 µg/mL) treatment.

| Species | Number of False Positives without PK* | Number of False Positives with PK* |
| --- | --- | --- |
| *E. coli* | 3/12 | 0/12 |
| *Klebsiella pneumoniae* | 0/7 | 0/7 |
| *Klebsiella oxytoca* | 0/7 | 0/7 |
| *Proteus mirabilis* | 0/3 | 0/3 |

\* A false positive was defined as a detection score greater than 1.5.

With the final workflow established for testing clinical isolates, we expanded on the earlier inhibitor experiments conducted with model laboratory CPO strains. Incorporating clinically employed carbapenemase inhibitors into the assay workflow would allow microbiology labs to rapidly characterize the inhibitor susceptibility profile of a particular CPO. Currently, the susceptibility of a CPO to β-lactam-β-lactamase inhibitor combinations is commonly tested by broth microdilution assays and E-tests, which suffer from long turnaround times (18 - 24 h).^29,30^ Recently, an extension panel for the automated VITEK2 antimicrobial susceptibility instrument was released, which includes the β-lactam-β-lactamase inhibitor combination of ceftazidime-avibactam. However, this panel is purchased separately from the base VITEK susceptibility card, thus increasing costs associated with testing. Incorporation of avibactam (4 µg/mL) into the biosensor assay workflow allowed for the successful differentiation between clinical CPOs which produce carbapenemases inhibited by avibactam (KPC, OXA-48-like) and those which carbapenemases that are not inhibited (NDM, VIM) (Figure 5). This information may prove useful in guiding treatment options for infections caused by multi-drug or pan-drug-resistant CPOs where therapeutic options are severely limited. The ability to rapidly determine the effectiveness of carbapenemase inhibitors against CPO isolates will gain importance with the continuing emergence of new carbapenemase variants which are less susceptible to currently used inhibitors.^26–28^

**Figure 5:**
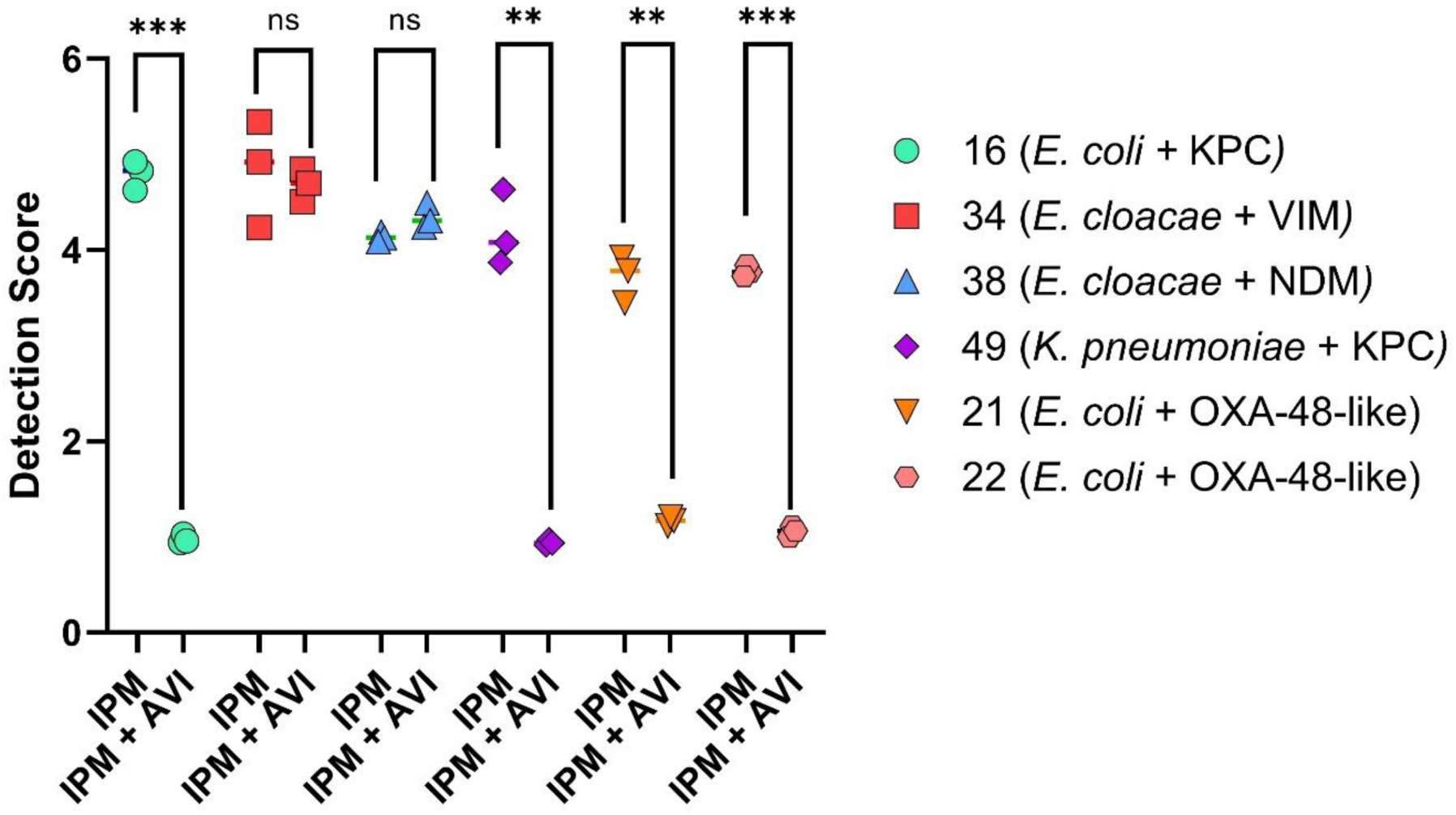
C**o**mparison **of detection scores between samples treated with imipenem (IPM) and avibactam (AVI), vs. imipenem alone.** A reduction in detection score for imipenem + avibactam- treated samples compared to imipenem alone is indicative of successful inhibition of the carbapenemase by avibactam. All isolates were tested in triplicate. ns = non significant; **: p < 0.01 ***: p < 0.001. Isolate numbers are indicated in the legend (Table S2).

Herein, we report the application of a luminescent whole-cell biosensor to the rapid detection of CPOs. The final optimized assay provides excellent results (100 % sensitivity, 100 % specificity) in approx. 2.5 hours inclusive of setup, thus making it a potentially useful option for rapid CPO screening. In addition to providing a binary response (yes/no) regarding whether a clinical isolate produces a carbapenemase, β-lactamase inhibitors such as avibactam can be easily introduced to the assay workflow to profile the inhibitor susceptibility of the carbapenemases(s) produced by a CPO.

One potential limitation of this assay relates to the measurement of the luminescent signal generated by the biosensor. In this study, we employed a multimode plate reader for measuring luminescence production by the biosensor, but such facilities might not be available in lower resource settings. Moving forward, the use of portable, point-of-care tube luminometers could serve as a less expensive option for the implementation of the biosensor assay. Furthermore, once a luminometer has been acquired, the cost-per-test is low (estimated to be approx. $ 0.47 USD per sample). To address limitations related to the need for a luminometer, colourimetric versions of this assay could be developed, thus allowing assay results to be interpreted visually without the need for instrumentation.

With the increasing prevalence of CPOs globally, rapid and accurate detection of these organisms will become even more imperative to ensure that critically ill patients receive timely and appropriate antimicrobial therapy. Additionally, specific infection prevention and control (IPAC) protocols (*e.g.,* contact precautions, patient isolation, increased room sanitation) are needed for patients who test positive for CPO infection or colonization.^31,47^ The use of rapid CPO screening methods will ensure that IPAC measures can be implemented rapidly in the case of positive results, thus aiding in the prevention of outbreaks in hospital wards and allowing for earlier de-escalation of these measures following a negative result. Outside of hospitals, surveillance of CPO prevalence in environmental and agricultural settings will continue to gain importance as the abundance of these organisms continues to increase. Thus, novel diagnostics for the rapid detection of CPOs such as that described here will help reduce the impact of these multi-drug-resistant pathogens on human health.

## MATERIALS AND METHODS

### Reagents

BD BBL cation-adjusted Mueller-Hinton Broth (CAMHB), bacterial protein extraction reagent (B-PER), proteinase K, Dulbecco’s phosphate buffered saline (DPBS) and phenol red were purchased from Fisher Scientific. Imipenem was purchased from Glentham Life Sciences. Avibactam was purchased from Medkoo Biosciences. BugBuster protein extraction reagent was purchased from MilliporeSigma. Nitrocefin was purchased from TOKU-E.

### Bacterial Strains

*E. coli* BW25113 (National BioResource Project, NBRP) was transformed with previously prepared pACYC184 vectors encoding representative penicillinases (TEM-116), ESBLs (CTX-M-15) and carbapenemases (NDM-1, IMP-1, KPC-2, OXA-48).^25^ The native promoter and gene encoding for OXA-10 was amplified from *Klebsiella pneumoniae* AR0041 by polymerase chain reaction (PCR) and cloned into the HindIII site of pACYC184 by HiFi assembly. The sequence of the recombinant plasmid was confirmed by Sanger sequencing (The Centre for Applied Genomics, The Hospital for Sick Children, Toronto, ON) and whole plasmid sequencing (Plasmidsaurus).

CPO clinical isolates were collected by the microbiology lab at Kingston Health Sciences Center (KHSC; Kingston, Canada) and ESBL-producing clinical isolates were collected by the Shared Hospital Laboratory (SHL; Toronto, Canada), all during the normal course of care. The panel of isolates consisted of 50 CPOs belonging to several different bacterial species (Table S2), and 30 ESBL producers (Table S3). The CPOs had been previously tested by the KHSC microbiology lab for carbapenemase production via the mCIM test, and the presence of carbapenemase genes (NDM, KPC, OXA-48, VIM) was confirmed by multiplex PCR. The presence of ESBLs was previously verified by SHL via the ESBL double disk synergy test; all of the ESBL-producing isolates tested negative for carbapenemase production based on meropenem and ertapenem minimum inhibitory concentrations (MICs).

All bacterial strains used in this study were cultured for 16 – 20 h at 37 °C on CAMHB agar plates. Carbapenemase-negative controls (*E. coli* BW25113, *E. coli* ATCC 25922) were grown on CAMHB plates lacking antibiotic. Laboratory *E. coli* BW25113 strains producing β- lactamases, clinical CPOs, and ESBL-producing isolates were cultured on CAMHB plates supplemented with either 25 µg/mL chloramphenicol, 1 µg/mL imipenem, or 100 µg/mL ampicillin, respectively. *E. coli* BW25113 cells transformed with the biosensor plasmid pAMPLUX ^25^ were cultured on CAMHB plates supplemented with 50 µg/mL kanamycin.

### Minimum Inhibitory Concentration Testing

All MIC testing was conducted according to CLSI guidelines.^48^ Test isolates were cultured as described above. Imipenem stocks were prepared in sterile water and serially diluted in CAMHB in clear sterile 96-well non-treated flat-bottom plates (Corning). Cell suspensions were prepared by suspending colonies in CAMHB to OD600 = 0.1 (approx. 1 x 10^8^ CFU/mL). These suspensions were diluted in CAMHB to approx. 5 x 10^6^ CFU/mL, and 20 µL of the diluted suspensions was added to 180 µL of each of the imipenem dilutions. Plates were incubated for 20 h at 37 °C without shaking and OD600 readings were collected using a Synergy LX plate reader (Agilent BioTek) to determine MICs.

### Laboratory Strain Carbapenemase Detection Assay

Test strains, a carbapenemase-negative control (*E. coli* BW25113) and biosensor cells were cultured as described above. Imipenem stocks were prepared in sterile water and diluted in the assay medium (CAMHB) to a final concentration of 1 µg/mL. In cases where avibactam was tested, avibactam stocks were prepared to 1 mg/mL in sterile water and diluted to 4 µg/mL in the assay medium (CAMHB) supplemented with 1 µg/mL imipenem.

A 1 µL loopful of the test strain or negative control was transferred to a 5 mL solution of imipenem (1 µg/mL) in CAMHB. In cases where avibactam was tested, a loopful of test strain or negative control was also transferred to a second tube containing a 5 mL solution of imipenem and avibactam (1 µg/mL and 4 µg/mL, respectively) in CAMHB. After mixing, 150 µL of this test solution was immediately transferred to 50 µL of a biosensor cell suspension [prepared by suspending biosensor cell colonies in CAMHB to an OD600 = 0.12, approximately 0.5 McFarland units (MFU)] in the wells of a white Lumitrac 200 96-well plate (Greiner Bio-One). Test mixtures were incubated at 37 °C without shaking for 90 minutes, and luminescence readings were taken using a Synergy LX multimode plate reader (Agilent BioTek). Detection scores were determined by dividing the luminescent readings for a carbapenemase-negative strain with the luminescent readings for the test strains. In experiments where avibactam was tested, detection scores were compared between samples treated with imipenem/avibactam vs. imipenem alone. All lab strains were tested in quadruplicate.

### Cell Lysis Testing

*E. coli* BW25113 transformed with pACYC184-NDM-1 was cultured as described above.

The following day, serial dilutions of BugBuster protein extraction reagent were prepared in autoclaved water to 100, 50, 25 and 0 % v/v. Then, 50 µL of each dilution was transferred to a 1.5 mL microcentrifuge tube. A 1 µL loopful of the NDM-1-producing *E. coli* was transferred to these lysis solutions, which were incubated at room temperature for 10 min. Cellular debris was pelleted for 3 min using a Fisherbrand 100-240V Mini-Centrifuge (fixed speed of 6,000 rpm). Following centrifugation, 10 µL of each supernatant was diluted in 990 µL of CAMHB, and 90 µL of the resulting solution was added to the wells of a sterile, clear, non-treated 96-well plate (Corning). Then, 10 µL of a 2 mM nitrocefin stock was added to the microplate wells, and reactions were monitored at 488 nm for 10 min using a Synergy LX multimode plate reader (Agilent BioTek).

### Initial Method for Carbapenemase Detection in Clinical Isolates

Test strains, a carbapenemase-negative control (*E. coli* ATCC 25922), and biosensor cells were cultured as described above. Imipenem stocks were prepared in sterile water, which were diluted to 1 µg/mL in CAMHB. Two 1 µL loopfuls of the test strains or control were added to 50 µL of 30% v/v BugBuster solution and incubated at room temperature for 10 min. Cellular debris was pelleted for 3 min using a Fisherbrand 100-240V Mini-Centrifuge (fixed speed of 6,000 rpm).

Following lysis, 10 µL of lysate was added to 990 µL of CAMHB containing 1 µg/mL imipenem, and the mixture was incubated for 30 min at 37 °C to allow for imipenem hydrolysis to occur. Following this incubation, 150 µL of the resulting solution was mixed with 50 µL of a 0.5 MFU suspension (in CAMHB) of biosensor cells in the wells of a white Lumitrac 200 96- well plate (Greiner Bio-One). These test mixtures were incubated at 37 °C without shaking for 90 min, and luminescence readings were taken using a Synergy LX multimode plate reader (Agilent BioTek). Detection scores were determined by dividing the luminescent readings for a carbapenemase-negative strain by the luminescent readings for the test strains. All isolates were tested in triplicate. Samples with a detection score greater than 1.5 were defined as positive; this value was chosen as it lies well above the average error on luminescence readings (approx. 5- 10 %).

### CARBA-NP Test

The CARBA-NP test was performed as outlined in the CLSI M100 Edition 34 document.^49^ CARBA-NP solution A was prepared by adding 2 mL of 0.5 % phenol red and 180 µL of 10 mM zinc sulfate to 16.6 mL of sterile water, which was then adjusted to pH 7.8. This solution was stored at 4 °C until the following day. Test strains and a carbapenemase-negative control (*E. coli*

ATCC 25922) were cultured as described above. CARBA-NP solution B was prepared on the day of the assay by dissolving imipenem in CARBA-NP solution A to a concentration of 3 mg/mL. A 1 µL loopful of each test strain was added to two separate 1.5 mL tubes containing 100 µL of B-PER and vortexed for 15 seconds. To the first set of tubes, 100 µL of solution A was added; to the second set of tubes, 100 µL of solution B was added. These mixtures were incubated at 37 °C for 2 h, and colour change was monitored after this incubation period.

### UV-Vis Spectrophotometric Imipenem Hydrolysis Assays

Test strains, a carbapenemase-negative control (*E. coli* ATCC 25922) and a carbapenemase-positive control (*E. coli* BW25113 pACYC184-OXA-48) were cultured as described above. Imipenem stocks were prepared in sterile water. A 10 µL loopful of each strain was added to 500 µL of 30 % v/v BugBuster solution and incubated at room temperature for 10 min. Cellular debris was pelleted for 10 min using a Fisherbrand 100-240V Mini-Centrifuge (fixed speed of 6,000 rpm). Supernatants were diluted 10X in DPBS lacking calcium and magnesium chloride, and 100 µL of the resulting solutions were added to the wells of a UV-Star 96-well plate (Greiner Bio-One) in triplicate. Next, 100 µL of a 60 µg/mL imipenem solution was added, and absorbance measurements at 297 nm were taken every minute for 10 min at room temperature using a Synergy LX multimode plate reader (Agilent BioTek). All samples were tested in triplicate.

### Impact of Strains Suspected of Producing Antibacterial Agents on Biosensor Luminescence

Test strains, a negative control (*E. coli* ATCC 25922) and biosensor cells were cultured as described above. Two 1 µL loopfuls of the test strains and negative control were added to 50 µL of 30 % v/v BugBuster solution and incubated at room temperature for 10 min. Cellular debris was pelleted for 3 min using a Fisherbrand 100-240V Mini-Centrifuge (fixed speed of 6,000 rpm). Following lysis, 10 µL of lysate was added to 990 µL of CAMHB. A 150 µL aliquot of this solution was mixed with 50 µL of a 0.5 MFU suspension of biosensor cells in the wells of a white Lumitrac 200 96-well plate (Greiner Bio-One). Luminescence readings were taken 0, 1 and 2 h post-treatment, with the plate incubated at 37 °C between readings. All samples were tested in triplicate.

### Carbapenemase Detection Assays Incorporating Proteinase K

Test strains, a carbapenemase-negative control (*E. coli* ATCC 25922), and biosensor cells were cultured as described above. Imipenem stocks were prepared in sterile water, then diluted to 1 µg/mL in CAMHB. Where applicable, avibactam stocks were prepared in sterile water and diluted to 4 µg/mL in CAMHB supplemented with imipenem. Proteinase K (PK) stocks were prepared to 20 mg/mL in sterile water, and aliquots were stored at -20 °C until use. Two 1 µL loopfuls of the test strains or negative control were added to 50 µL of 30% v/v BugBuster solution and incubated at room temperature for 10 min. Cellular debris was pelleted for 3 min using a Fisherbrand 100-240V Mini-Centrifuge (fixed speed of 6,000 rpm). Following lysis, 10 µL of lysate was added to 990 µL of CAMHB supplemented with 1 µg/mL imipenem. In cases where avibactam was tested, 10 µL of lysate was added to a second tube containing 990 µL CAMHB solution with imipenem and avibactam (1 µg/mL and 4 µg/mL, respectively). The mixtures were incubated for 30 min at 37 °C. Following this incubation period, the PK stock solution was added to each tube to a final concentration of 100 µg/mL, which were then incubated at 37 °C for 15 min. Following this incubation, 150 µL of the resulting solution was mixed with 50 µL of a 0.5 MFU suspension (in CAMHB) of biosensor cells in the wells of a white Lumitrac 200 96-well plate (Greiner Bio-One). These test mixtures were incubated at 37 °C without shaking for 90 min, and luminescence readings were taken using a Synergy LX multimode plate reader (Agilent BioTek). Detection scores were determined by dividing the luminescent readings for a carbapenemase-negative sample by the luminescent readings for the test samples. All isolates were tested in triplicate.

### Statistical Analysis

All statistical tests were conducted using GraphPad Prism, version 10.1. In all cases, unpaired t-tests with Welch’s correction were used to determine statistical significance between detection scores for relevant treatment populations.

## AUTHOR INFORMATION

Corresponding Author Christopher T. Lohans

## Acknowledgements

The authors wish to thank the clinical microbiology laboratory staff at Kingston Health Sciences Center (Kingston, Ontario) and Shared Hospital Labs (Toronto, Ontario) for providing bacterial isolates and phenotypic and genotypic information to aid this study.

## Author Contributions

M.A.J. designed the study, prepared the manuscript, and conducted the majority of the experiments. S.B., J.L.L., and R.A.V.G. contributed to the laboratory experiments. X.X.L. and P.M.S. provided clinical isolates. C.T.L. designed the study and prepared the manuscript. All authors approved the submitted manuscript.

## ABBREVIATIONS

AVI, avibactam; B-PER, bacterial protein extraction reagent; CAMHB, cation-adjusted Mueller-Hinton broth; CLSI, Clinical and Laboratory Standards Institute; CPO, carbapenemase-producing organism; DPBS, Dulbecco’s phosphate buffered saline; ESBL, extended spectrum beta-lactamase; GES, Guiana extended-spectrum beta-lactamase; IPAC, infection prevention and control; KHSC, Kingston Health Sciences Center; KPC, *Klebsiella pneumoniae* carbapenemase; IMP, imipenemase; MBL, metallo-beta-lactamase; MFU, McFarland units; mCIM, modified carbapenem inactivation method; MIC, minimum inhibitory concentration; NDM, New Delhi metallo-beta-lactamase; OXA, oxacillinase; PBP, penicillin-binding protein; PCR, polymerase chain reaction; PK, proteinase K; SHL, Shared Hospital Lab; VIM, Verona integron-encoded metallo-beta-lactamase.

