## Supplemental Information for "Rapid Detection of Carbapenemase-Producing Organisms Using a Luminescent Biosensor"

### SI Tables and Figures:

**Table S1. Imipenem minimum inhibitory concentrations (MICs) of the *Escherichia coli* laboratory strains used in this study.**

| ***E. coli*** **strain** | **Imipenem MIC (μg/mL)** |
|---|---|
| ATCC 25922 | 0.25 |
| BW25113 WT | 0.25 |
| BW25113 Δ*ompF* | 0.5 |
| BW25113 + pACYC184-NDM-1 | 512 |
| BW25113 + pACYC184-IMP-1 | 256 |
| BW25113 + pACYC184-KPC-2 | 256 |
| BW25113 + pACYC184-OXA-48 | 4 |
| BW25113 + pACYC184-CTX-M-15 | 0.25 |
| BW25113 + pACYC184-OXA-10 | 0.25 |
| BW25113 + pACYC184-TEM-116 | 0.25 |

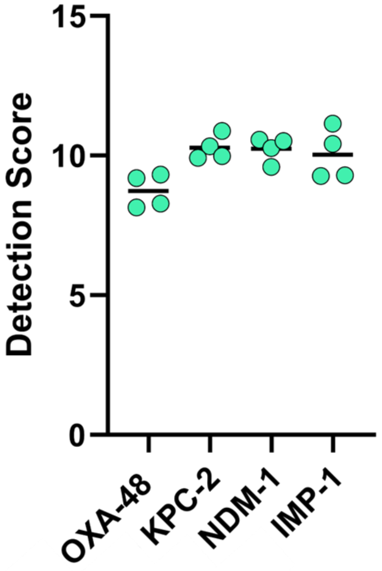

**Figure S1: Validation of the biosensor assay with carbapenemase-producing *E. coli* Δ*ompF* strains.** Detection scores for *E. coli* BW25113 Δ*ompF* cells expressing the carbapenemases OXA-48, NDM-1, IMP-1, and KPC-2. Detection scores were determined by dividing the luminescent readings for a carbapenemase-negative sample with the luminescent readings for the test samples. All strains were tested in quadruplicate.

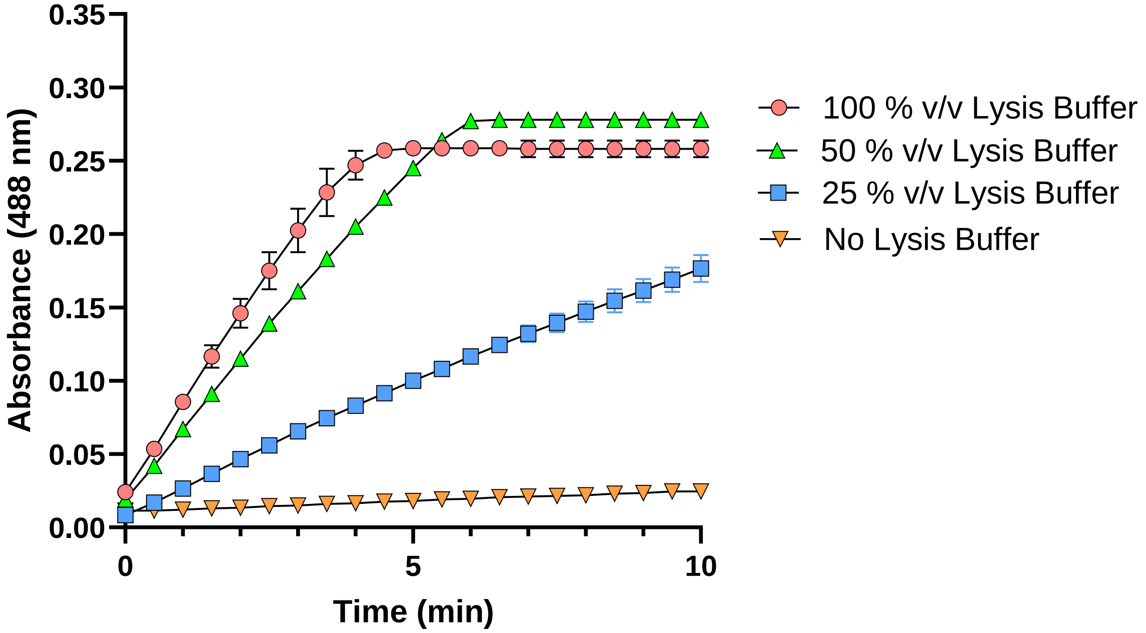

**Figure S2: Impact of different BugBuster dilutions on the release of NDM-1 from *E. coli* BW25113 cells.** NDM-1-producing *E. coli* were subjected to various BugBuster (lysis buffer) treatments ranging from 0 % v/v to 100 % v/v. The rate of nitrocefin hydrolysis (measured at 488 nm) was used as a surrogate marker for the release of NDM-1 from *E. coli* cells. Treatment of NDM-1-producing *E. coli* with 100, 50 and 25 % BugBuster increased the rate of nitrocefin hydrolysis compared to unlysed samples. All conditions were tested in triplicate, and error bars indicate S.D.

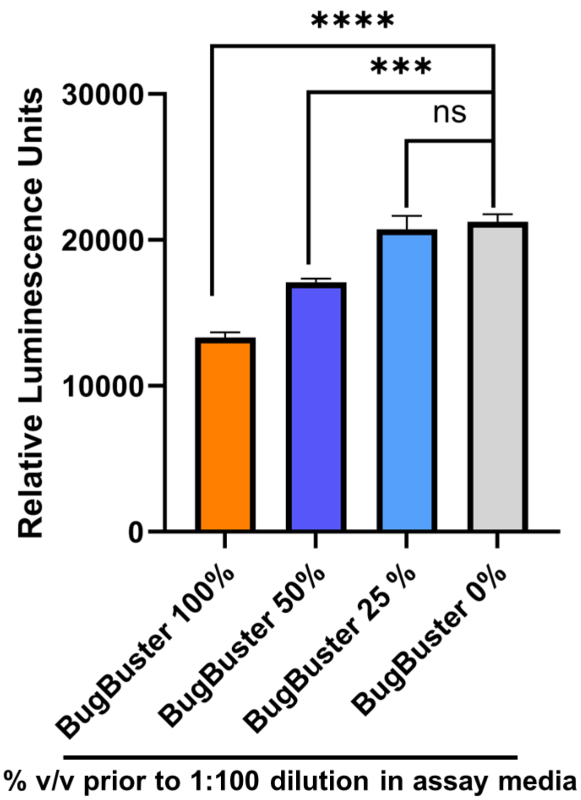

**Figure S3: Impact of BugBuster dilutions on luminescence production by biosensor cells.** Biosensor cells (*E. coli* BW25113 + pAMPLUX) were treated with assay media (CAMHB + 1 µg/mL imipenem) supplemented with varying concentrations of BugBuster for 1.5 h. BugBuster solutions were prepared to 100 %, 50 % and 25 % v/v prior to being further diluted 1:100 in the assay media. All conditions were tested in triplicate, and error bars indicate S.D. ns = nonsignificant; ***: $p < 0.001$; ****: $p < 0.0001$.

**Table S2. Imipenem MICs, biosensor detection scores, and biosensor test results for the clinical CPO isolates used in this study**.

| ISOLATE # | Species | Enzyme | Imipenem MIC (µg/mL) | Detection Score | Biosensor Result* |
|---|---|---|---|---|---|
| **1** | *Citrobacter freundii* | VIM | 8 | 5.8 | Positive |
| **2** | *C. freundii* | VIM | 16 | 5.7 | Positive |
| **3** | *C. freundii* | KPC | 8 | 6.6 | Positive |
| **4** | *C. freundii* | KPC | 4 | 6.6 | Positive |
| **5** | *C. freundii* | KPC | 8 | 6.3 | Positive |
| **6** | *C. freundii* | KPC | 8 | 6.7 | Positive |
| **7** | *C. freundii* | KPC + NDM | 16 | 6.2 | Positive |
| **8** | *E. coli* | VIM | 8 | 6.6 | Positive |
| **9** | *E. coli* | NDM | 16 | 6.0 | Positive |
| **10** | *E. coli* | NDM | 8 | 5.9 | Positive |
| **11** | *E. coli* | NDM | 16 | 5.2 | Positive |
| **12** | *E. coli* | NDM | > 32 | 5.5 | Positive |
| **13** | *E. coli* | NDM | 8 | 6.7 | Positive |
| **14** | *E. coli* | NDM | 8 | 6.4 | Positive |
| **15** | *E. coli* | NDM | > 32 | 6.6 | Positive |
| **16** | *E. coli* | KPC | 4 | 6.2 | Positive |
| **17** | *E. coli* | KPC | 2 | 4.8 | Positive |
| **18** | *E. coli* | KPC | 4 | 5.5 | Positive |
| **19** | *E. coli* | KPC | 4 | 6.5 | Positive |
| **20** | *E. coli* | KPC | 8 | 5.4 | Positive |
| **21** | *E. coli* | OXA-48-like | 2 | 3.5 | Positive |
| **22** | *E. coli* | OXA-48-like | 2 | 3.9 | Positive |
| **23** | *E. coli* | OXA-48-like | 32 | 6.2 | Positive |
| **24** | *E. coli* | OXA-48-like | 16 | 5.1 | Positive |
| **25** | *E. coli* | OXA-48-like | 16 | 5.4 | Positive |
| **26** | *E. coli* | OXA-48-like | 2 | 5.2 | Positive |
| **27** | *E. coli* | OXA-48-like | 16 | 5.1 | Positive |
| **28** | *E. coli* | OXA-48-like | 32 | 4.7 | Positive |
| **29** | *Enterobacter cloacae* | VIM | 32 | 4.9 | Positive |
| **30** | *E. cloacae* | VIM | > 32 | 8.4 | Positive |

| | | | | | |
|---|---|---|---|---|---|
| **31** | *E. cloacae* | VIM | 16 | 5.2 | Positive |
| **32** | *E. cloacae* | VIM | 32 | 5.9 | Positive |
| **33** | *E. cloacae* | VIM | 16 | 6.0 | Positive |
| **34** | *E. cloacae* | VIM | 4 | 6.2 | Positive |
| **35** | *E. cloacae* | VIM | 4 | 6.4 | Positive |
| **36** | *E. cloacae* | VIM | 16 | 7.1 | Positive |
| **37** | *E. cloacae* | VIM | 16 | 7.4 | Positive |
| **38** | *E. cloacae* | NDM | 8 | 4.8 | Positive |
| **39** | *E. cloacae* | NDM | 16 | 6.3 | Positive |
| **40** | *E. cloacae* | NDM | > 32 | 6.8 | Positive |
| **41** | *Klebsiella oxytoca* | KPC | 32 | 6.5 | Positive |
| **42** | *Klebsiella pneumoniae* | VIM | 4 | 8.8 | Positive |
| **43** | *K. pneumoniae* | NDM | 32 | 6.4 | Positive |
| **44** | *K. pneumoniae* | NDM | 4 | 5.5 | Positive |
| **45** | *K. pneumoniae* | KPC | 8 | 6.3 | Positive |
| **46** | *K. pneumoniae* | KPC | 4 | 7.1 | Positive |
| **47** | *K. pneumoniae* | KPC | 4 | 6.8 | Positive |
| **48** | *K. pneumoniae* | KPC | 8 | 5.9 | Positive |
| **49** | *K. pneumoniae* | KPC | 4 | 6.9 | Positive |
| **50** | *Morganella morganii* | VIM | 8 | 5.9 | Positive |

*** Samples which yielded detection scores greater than 1.5 were defined as positive**

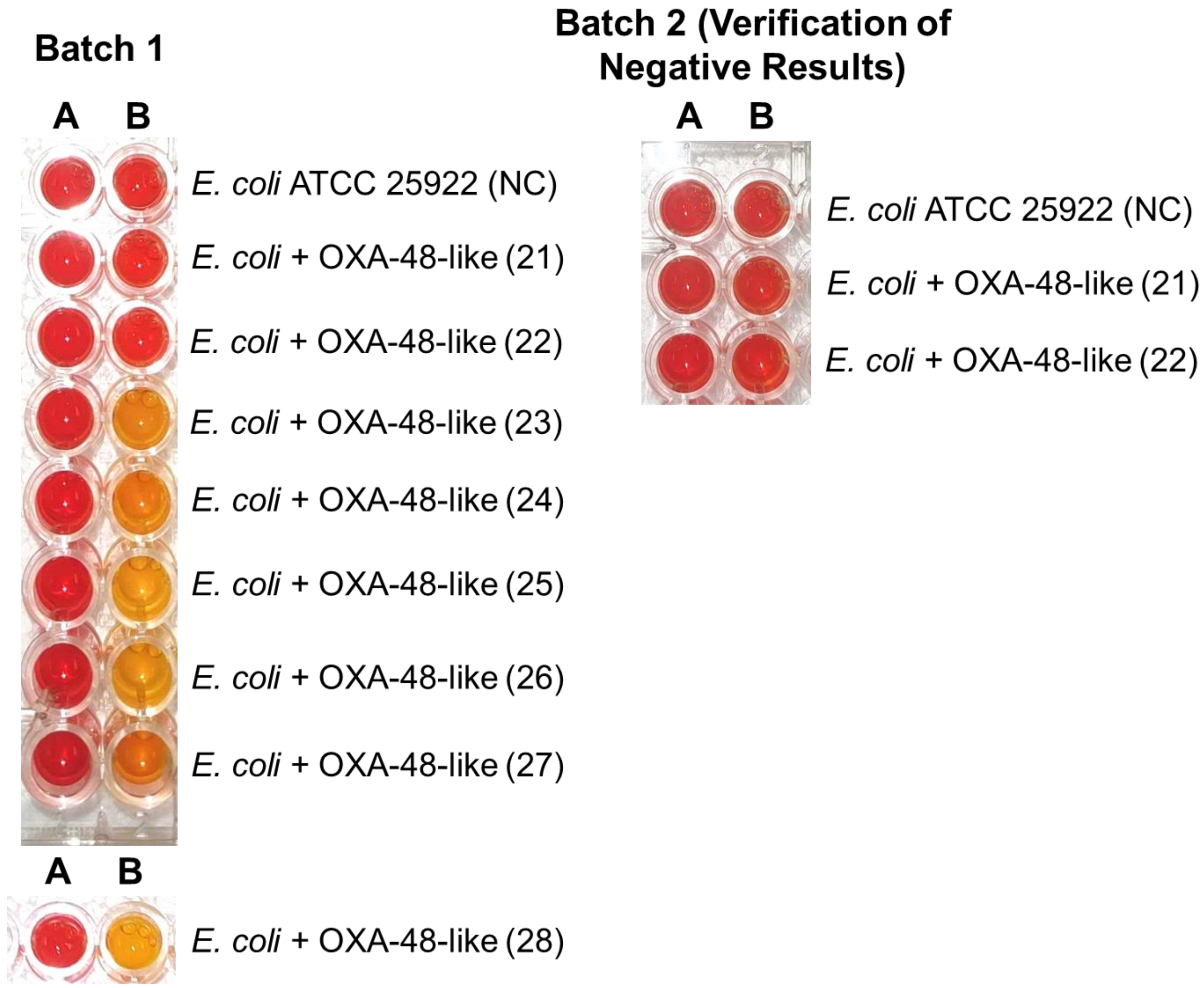

**Figure S4: CARBA-NP results for OXA-48-like-producing *E. coli* clinical isolates.** The labels above the columns of the microplate indicate whether samples were treated with CARBA-NP solution A or solution B (*i.e.*, solution A + 3 mg/mL imipenem). All OXA-48-like-producers were tested in batch 1, while batch 2 verified the negative results obtained for isolates 21 and 22 in a second, independent experiment. As per CLSI interpretation guidelines, a colour change from red to yellow indicates a sample is positive for carbapenemase, while no colour change (*i.e.*, red, dark orange) is considered a negative result. Samples with insufficient colour change (*i.e.,* orange) are considered invalid. Based on these criteria, six isolates tested positive and two isolates tested negative. Interpretation criteria were obtained from the CLSI M100 34$^{th}$ edition.[1]

**Table S3. Detection scores and biosensor test results [with and without proteinase K (PK) treatment] for the clinical non-CPO isolates used in this study**.

| ISOLATE # | Species | ESBL Class* | Detection Score (no PK) | Biosensor Result (no PK)*** | Detection Score (+ PK) | Biosensor Result (+ PK)*** |
|---|---|---|---|---|---|---|
| **N1** | *E. coli* | D | 1.0 | Negative | 1.0 | Negative |
| **N2** | *E. coli* | C | 1.0 | Negative | 0.9 | Negative |
| **N3**** | *K. pneumoniae* | A | ------------ | ------------- | ------------ | ------------- |
| **N4** | *E. coli* | C | 5.1 | **Positive** | 0.9 | **Negative** |
| **N5** | *K. oxytoca* | A | 0.9 | Negative | 1.0 | Negative |
| **N6** | *K. oxytoca* | A | 1.0 | Negative | 1.0 | Negative |
| **N7** | *K. pneumoniae* | A | 1.0 | Negative | 1.0 | Negative |
| **N8** | *E. coli* | A | 1.1 | Negative | 1.1 | Negative |
| **N9** | *E. coli* | A | 1.0 | Negative | 1.0 | Negative |
| **N10** | *E. coli* | A + C | 1.0 | Negative | 1.0 | Negative |
| **N11** | *K. pneumoniae* | C | 1.0 | Negative | 1.0 | Negative |
| **N12** | *E. coli* | A + C | 1.1 | Negative | 1.0 | Negative |
| **N13** | *E. coli* | A + C | 1.0 | Negative | 0.9 | Negative |
| **N14** | *K. pneumoniae* | A + C | 1.0 | Negative | 0.9 | Negative |
| **N15** | *K. pneumoniae* | A + C | 1.0 | Negative | 0.9 | Negative |
| **N16** | *Proteus mirabilis* | A | 1.0 | Negative | 0.9 | Negative |
| **N17** | *P. mirabilis* | A | 1.0 | Negative | 0.9 | Negative |
| **N18** | *K. oxytoca* | A | 1.0 | Negative | 0.9 | Negative |
| **N19** | *K. oxytoca* | D | 0.9 | Negative | 0.8 | Negative |
| **N20** | *P. mirabilis* | A | 1.1 | Negative | 1.0 | Negative |
| **N21** | *E. coli* | C | 1.0 | Negative | 0.9 | Negative |
| **N22** | *E. coli* | C | 1.0 | Negative | 1.0 | Negative |
| **N23** | *E. coli* | C | 27.9 | **Positive** | 0.9 | **Negative** |
| **N24** | *E. coli* | C | 31.3 | **Positive** | 0.9 | **Negative** |
| **N25** | *K. pneumoniae* | A | 1.0 | Negative | 1.0 | Negative |
| **N26** | *K. pneumoniae* | A | 1.0 | Negative | 0.9 | Negative |
| **N27** | *K. pneumoniae* | A | 0.9 | Negative | 0.9 | Negative |
| **N28** | *K. oxytoca* | A | 1.1 | Negative | 1.0 | Negative |
| **N29** | *K. oxytoca* | A | 1.0 | Negative | 1.1 | Negative |
| **N30** | *K. oxytoca* | A | 1.0 | Negative | 1.0 | Negative |

***The presence of ESBLs was verified by via the ESBL double disk synergy test**

****Repeatedly observed insufficient growth of this isolate on CAMHB-ampicillin agar plates for testing.**

*****Samples which yielded detection scores greater than 1.5 were defined as positive, while samples with scores less than 1.5 were defined as negative**

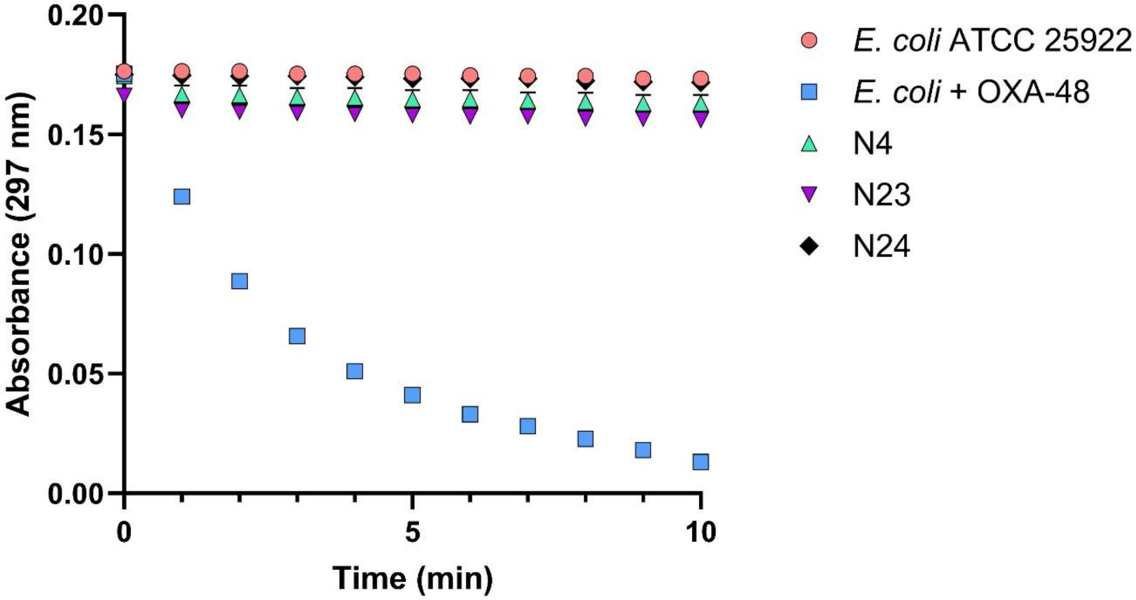

**Figure S5: UV-Vis spectrophotometric assay measuring imipenem hydrolysis by non-carbapenemase-producing *E. coli* ATCC 25922, OXA-48-producing *E. coli* BW25113, and test strains N4, N23, and N24.** Imipenem hydrolysis by diluted cell lysates was measured by monitoring the absorbance of intact imipenem at 297 nm. Reduction of absorbance is indicative of imipenem hydrolysis. All strains were tested in triplicate, and error bars indicate S.D.

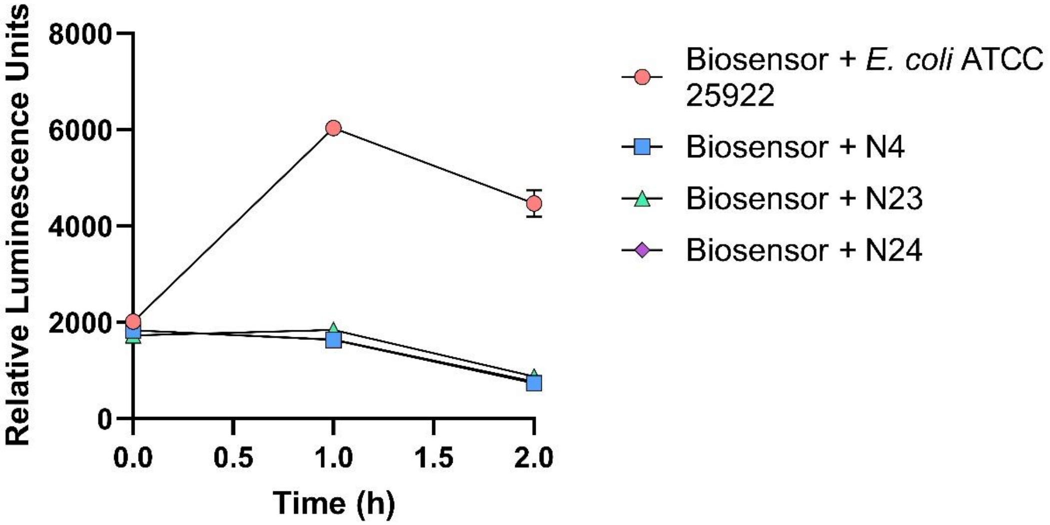

**Figure S6: Impact of lysates of *E. coli* ATCC 25922 and test isolates N4, N23 and N24 on biosensor luminescence in the absence of imipenem**. Luminescence production by *E. coli* biosensor cells was measured 0, 1 and 2 h post treatment with 1:100-diluted cell lysates. Exposure of biosensor cells to lysates from test strains N4, N23 and N24 resulted in reduced luminescence production after 1 and 2 h, relative to the lysate of the *E. coli* ATCC 25922 control. All strains were tested in triplicate, and error bars indicate S.D.

**Table S4:** Detection scores for representative CPO isolates tested with and without PK (100 µg/mL) treatment.

| **Isolate*** | **Detection Score (no PK)** | **Detection Score (+ PK)** |
|---|---|---|
| 21 (*E. coli* + OXA-48-like) | 3.3 | 3.2 |
| 30 (*E. cloacae* + VIM) | 8.4 | 5.6 |
| 43 (*K. pneumoniae* + NDM) | 5.1 | 4.7 |
| 45 (*K. pneumoniae* + KPC) | 5.6 | 5.7 |

### References

(1) Clinical and Laboratory Standards Institute. *CLSI M100 Performance Standards for Antimicrobial Susceptibility Testing 34th Edition.*; 2024.
